# Rounding up the annual ryegrass genome: high-quality reference genome of *Lolium rigidum*

**DOI:** 10.1101/2022.07.18.499821

**Authors:** Jefferson Paril, Gunjan Pandey, Emma B. Barnett, Rahul V. Rane, Leon Court, Thomas Walsh, Alexandre Fournier-Level

**Affiliations:** School of BioSciences, University of Melbourne, Parkville, Australia; CSIRO Land and Water, Acton, Australia; CSIRO Health and Biosecurity, Parkville, Australia

## Abstract

The genome of the major agricultural weed species, annual ryegrass (*Lolium rigidum*) was assembled, annotated and analysed. Annual ryegrass is a major weed in wheat cropping, and has the remarkable capacity to evolve resistance to herbicides with various modes of action. The chromosome-level assembly was achieved using short- and long-read sequencing in combination with Hi-C mapping. The assembly size is 2.44Gb with N_50_=361.79Mb across 1,764 scaffolds where the seven longest sequences correspond to the seven chromosomes. Genome completeness assessed through BUSCO returned a 99.8% score for complete (unique and duplicated) and fragmented genes using the Viridiplantae set. We found evidence for the expansion of herbicide resistance-related gene families including detoxification genes. The reference genome assembly of *L. rigidum* is pivotal for the management of this highly problematic weed species which leverages genomic tools to devise new control options.

## Introduction

*Lolium rigidum* (Gaudin, 1811) also known as annual ryegrass, rigid ryegrass, or Wimmera grass, is the world’s most herbicide resistant weed species. It has developed resistance to over a dozen different modes of action across a number of herbicides and has the highest incidence of resistance in any weed species (Heap 2022). It is the first weed species reported to have evolved resistance to glyphosate (Powles et al. 1998).

*L. rigidum* is a diploid grass species with a chromosome number of 2n=2x=14 (Terrell 1966; Monaghan 1980) and an estimated genome size of ∼2Gb, similar to that of the closely-related forage crop *Lolium perenne* (Byrne et al. 2015; Frei et al, 2021). This species known to hybridise with other members of the *Lolium* genus, e.g. *L. multiflorum* and *L. perenne* (Kloot 1983). This genus is thus a complex of cross-compatible species which can produce fertile hybrids and makes species boundaries ambiguous (Naylor 1960; Terrell 1966; Kloot 1983).

*L. rigidum* is a highly-competitive, self-incompatible, wind-pollinated, annual, C3 weed species (Monaghan 1980; McCraw et al. 1983), which can produce up to 45,000 seeds per square metre in wheat fields where it can achieve high densities (33%-67% abundance in agricultural field conditions; Rerkasem et al, 1980). The combination of its high fecundity and outcrossing reproduction regime results in large and genetically diverse populations with high adaptive potential. The seeds have varying levels of dormancy ensuring its persistence in the soil seedbank (Goggin et al. 2012). Ryegrass infestation causes significant yield reduction in rapeseed and cereal crops (Lemerle et al. 1995) and its seeds can get infected with *Clavibacter toxicus* causing livestock poisoning (Riley and McKay 1991; Ophel et al. 1993).

*L. rigidum* is native to the Mediterranean region and was widely introduced around the world as a pasture crop. In the 19^th^ century, it was introduced to Australia (Kloot 1983) where it successfully adapted through a combination of artificial and natural selection. It is now the major weed in the wheat-growing regions of Australia (Reeves 1976; Medd et al. 1985; Powles and Matthews 1992).

In this paper, we report the first chromosomal-level genome assembly *Lolium rigidum*. This information is a valuable resource towards genomically-informed management of this major agricultural weed species with a particular emphasis on the issue of herbicide resistance evolution.

## Materials and Methods

### Plant sampling, tissue culture, and DNA extraction

A single glyphosate-resistant plant from Wagga Wagga (NSW, Australia) was selected as the reference genome for *Lolium rigidum*. This individual was tissue-cultured to induce embryogenic calli for clonal multiplication and maintenance following the protocol for *Lolium* spp. by Creemers-Molenaar and Beerepoot (1992). DNA was extracted using Qiagen DNeasy plant mini kit (QIAGEN N.V., Venlo, Netherlands) following manufacturer’s instructions.

### Genome sequencing and assembly

Short- and long-read DNA sequence data were generated and scaffolded using Hi-C sequence information. Short-read sequencing libraries were constructed using NEBNext Ultra II DNA Library Prep kit for Illumina (NEB, USA) at the University of Melbourne, Australia, and sequenced using HiSeq X platform (Illumina, Inc., San Diego, USA) ran in 150-bp paired-end mode (Azenta Life Sciences, Suzhou, China). Adapter sequences were removed from the resulting reads using TrimGalore (v 0.6.6). Long read sequencing was carried out on MinION and PromethION platforms. Basecalling was performed using *guppy* (v5.1; Wick et al, 2019) under the *dna_r9*.*4*.*1_450bps_sup*.*cfg* model. The long-read sequences were trimmed using *Porechop* (v0.2.4; Wick et al, 2017) and filtered using *filtlong* (v0.2.1) to obtain high quality reads. The long-reads were assembled using *Flye* (v2.9; Kolmogorov et al, 2020) with the minimum overlap parameter set to 6,000. Duplicate contigs were purged using *purge_dups* (v1.2.5; Guan et al, 2020) with the default settings. The long-reads were error-corrected and trimmed using *Canu* (v2.2; Koren et al, 2017), and were used in three rounds of contig polishing using *Racon* (v1.4.22; Vaser et al, 2017). This was followed by three rounds of short-read-based polishing using *Polca* (*MaSURCA* v4.0.7; Zimin et al, 2013) to obtain the final contig assembly. This assembly was assessed using *BUSCO* (v5; Simão et al, 2015) against the Viridiplantae and Poales lineages’ gene sets.

A Hi-C library was prepared using 20 mg of leaf tissue and the Arima HiC kit following the manufacturer’s instructions. The library was sequenced on Novaseq 6000 platform (Illumina, Inc., San Diego, USA) to generate 500 million reads. The final contig assembly was scaffolded based on the genomic topological information using *ALLHiC* (v1; Zhang et al, 2019) and polished using *3d-dna* (Dudchenko et al, 2017). Genome size was estimated based on the kmer distribution of the Illumina sequences using Jellyfish 2.3.0 (Marcais and Kingsford, 2011) at kmer=19.

The assembly is available on the National Center for Biotechnology Information of the United States (NCBI) database under the accession number (SAMN25144995, JAKKIG000000000). Raw Illumina, MinION, PromethION, and Hi-C reads are available under the NCBI Bioproject PRJNA799061.

### Transcriptome sequencing, assembly, and genome annotation

Clones from the reference plant established through tissue culture were grown under greenhouse conditions. Two independent samples of each whole seedlings, roots, stems, leaves, inflorescence, meristems were snap-frozen and total RNA was extracted using Isolate II RNA plant kit (Bioline, UK). RNA-sequencing libraries were synthesised for each sample using NEBNext Ultra II stranded RNA library synthesis kits, indexed using the NEBNext Multiplex Oligos for Illumina barcode kit. Libraries were quantified using NEBNext Library Quant KIt for Illumina, normalised, pooled, and sequenced on an Illumina HiSeq X ten platform at Azenta Life Sciences (Suzhou, China) to generate ∼257 million 150-bp paired-end reads. The reads were demultiplexed and error-corrected using Rcorrector (v1.0.4; Song et al, 2015). Adapters and low quality base pairs were trimmed using TrimGalore (v0.6.0). Ribosomal RNA sequences were discarded when one of the paired-end reads mapped to the sequences present in the SILVA database (v138.1; Quast et al, 2012) using Bowtie2 (v2.3; Langmead & Salzberg, 2012). After filtering, ∼197 million reads were used for *de novo* transcriptome assembly using the De novo RNA-Seq Assembly Pipeline (Cabau et al, 2017) including the rice protein sequences (release 51) as guide and using both Trinity (v2.8.4; Haas et al, 2013) and Oases (v0.2.09; Schulz et al, 2012) as assemblers. The resulting two assemblies were merged into a single compacted meta-assembly. The filtered reads were re-mapped against the meta-assembly and transcripts with FPKM>1 were included in the transcriptome.

The genome was annotated using NCBI’s genome annotation pipeline using the *de novo* assembled transcriptome. Transposable elements were identified using RepeatMasker and RepeatModeller (v4.1.2 and v2.0.3, respectively; Flynn et al, 2020).

### Comparative genomics

The reference genomes, annotations and coding DNA sequences (CDS) of *Arabidopsis thaliana* (TAIR10 v1), rice (*Oryza sativa*; IRGSP v1), sorghum (*Sorghum bicolor*; NCBI v3), maize (*Zea mays*; B73 Reference NAM v5), and perennial ryegrass (*Lolium perenne*; Kyuss v1) were used for comparative genomics analyses. The well curated genome of *A. thaliana* was used as the outgroup. Rice and maize genomes represent well annotated grass genomes. Perennial ryegrass is a closely related species. Sorghum is an additional grass crop species.

OrthoFinder (v2.5.4; Emms and Keyll 2018) were used to cluster the CDS of all six species into orthogroups, i.e. similar genes including paralogs within species and orthologs among species. The resulting orthogroups were assigned to gene families they most likely belong to using HMMER (v3.3.2; Mistry et al, 2013) and PantherHMM gene family models (v17; Mi et al, 2019). Significant gene family contraction and expansion in each of the six genomes were determined using CAFE (v5; De Bie et al, 2006) and a P-value<0.01. The significantly expanded gene families were used for gene ontology (GO) enrichment analysis using the GO consortium’s web tool “Gene ontology enrichment analysis tool” (The UniProt Consortium, 2019).

Orthogroups consisting of a single gene in each of the six genomes, i.e. single-copy gene orthogroups were used to generate a phylogenetic tree by maximum likelihood, and to estimate the pairwise sequence divergence times between species by empirical Bayesian approach. These single-copy gene orthogroups were aligned using MACSE (v2.06; Ranwez et al, 2011). The phylogenetic tree was generated using IQ-TREE (v2.0.7; Minh et al, 2020) and TimeTree.org fossil record estimates, i.e. median divergence times between:

- *A. thaliana* & rice: 160 million years ago (MYA),
- sorghum & perennial ryegrass: 62 MYA, and
- perennial ryegrass & annual ryegrass: 2.74 MYA.

The rate of transversions at four-fold degenerate sites (4DTv) for each sequence pair across paralogs within species and orthologs across species was calculated to estimate relative divergence times and identify whole genome duplication (WGD) events.

### Herbicide resistance genes

The use of a glyphosate-resistance plant as reference genome lends itself to the identification of the potential genomic basis of herbicide resistance. These resistance-conferring genomic features may be point mutations in genes coding for essential enzymes targeted by herbicides or in detoxification genes. These mutations can be detected using pairwise rates of synonymous and non-synonymous substitutions, i.e. Ka/Ks ratio (1: neutral, >1:positive selection, <1: stabilising selection), estimated between homologous pairs of protein coding sequences. Additionally, these resistance-conferring features may be structural changes leading to gene loss or duplication. These can be detected by assessing the patterns of expansion and contraction in the genes coding for the target of glyphosate (i.e. enolpyruvylshikimate phosphate synthase; EPSPS indispensable for aromatic amino acid synthesis), and detoxification-related gene families.

To perform these analyses, 8 herbicide resistance-related enzymes were used: enolpyruvylshikimate phosphate synthase (EPSPS), superoxide dismutase (SOD), ascorbate peroxidase (APX), glutathione S-transferase (GST), monodehydroascorbate reductase (MDAR), glutathione peroxidase (GPX), cytochrome P450 (CYP450), and ATP-binding cassette transporter (ABC). The sequences of these proteins were downloaded from the Universal Protein Resource (UniProt) database. Protein sequences specific to *L. rigidum, L. multiflorum, A. thaliana, O. sativa*, and *Z. mays* were queried because of the relatively high annotation qualities in these species. The predicted protein sequences of the six genomes were queried against the herbicide-resistance-related protein sequences and the best matching encoding gene was identified (E-value≤1×10^−10^). Significantly expanded and contracted TSR and NTSR gene families using all six species were identified using CAFE (v5; De Bie et al, 2006) and a P-value<0.01. The coding sequences of EPSPS gene paralogs within the annual ryegrass genome and homologs in the other 6 genomes were assessed using Ka/Ks ratio across 15-bp non-overlapping sliding windows using KaKs_calculator (version 2; Wang et al, 2009).

## Results

### Genome assembly

A total of 294.8 Gb from short- and long-read whole-genome sequencing was generated. Illumina sequencing of 150-bp paired-end reads accounted for 68.69% of this output (202.5 Gb across ∼1.35 billion reads). Oxford Nanopore sequencing using MinION and Promethion platforms accounted for 31.31% of the sequencing output (92.3 Gb across 9.04 million reads with N_50_=16.2kb). The Hi-C library generated 66 Gb of raw sequence data. We estimated the genome size to be 2.40 Gb based on k-mer frequency analysis (k=19).

The final assembly reached a chromosomal level with a total length 2.44Gb (N_50_=361.79Mb) over 1,764 scaffolds where the 7 longest sequences constituting 96% of the assembly correspond to the 7 chromosomes (Table 1 and Supplementary Figure 1). This assembly is 99.8% complete based on BUSCO analysis using the Viridiplantae gene set (i.e. Complete & single-copy [S]: 29.4%, Complete & duplicated [D]: 70.4%, Fragmented [F]: 0.2%, Missing [M]: 0.0%, Total [n]: 425) and 97.2% complete using the Poales gene set (S:32.4%, D:64.8%, F:0.9%, M:1.9%, n:4896).

**Table 1.** *Lolium rigidum* genome assembly (NCBI RefSeq: GCF_022539505.1) statistics.

|  |  |
| --- | --- |
| Estimated genome size | 2.40 Gbp |
| **Total assembly size** | 2.44 Gbp |
| **Total scaffolds** | 1,764 |
| **N<sub>50</sub>** | 0.36 Gbp |
| **L<sub>50</sub>** | 4 |
| **Longest scaffold** | 0.41 Gbp |
| **Number of chromosomes** | 7 |
| **Total chromosome length** | 2.35 Gbp |
| **GC content** | 44.76% |
| **Total length of retroelements** | 0.82 Gb (33.61%) |
| **Total length of DNA transposons** | 0.09 Gb (3.84%) |
| **Number of gene models** | 57,529 |
| **Mean gene length** | 3,624 bp |

**Figure 1.**
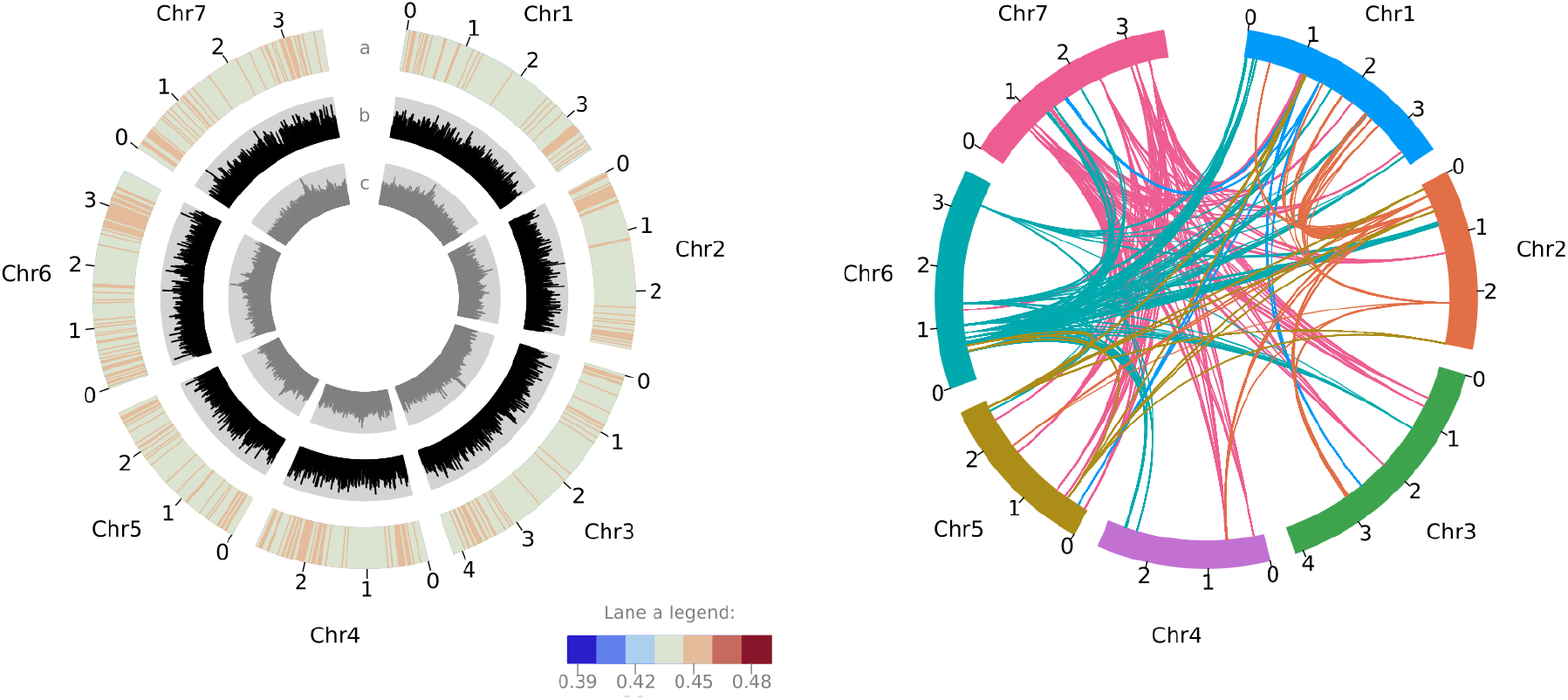
Features of the *Lolium rigidum* genome (each tick is ×100Mb). **Left panel - lane a**: GC content heatmap of mean GC content per 2.35Mb window (ranging from 42% to 47%); **lane b**: distribution of Copia long terminal repeat (LTR) retrotransposon family; **lane c**: distribution of Gypsy LTR retrotransposon family. **Right panel**: chord diagram shows the syntenic relationships within the top 5 orthogroups with the most paralogs in the genome, where the colours match the colours of the chromosome most of the paralogs per orthogroup are located.

Our *L. rigidum* genome assembly mainly consists of interspersed repeats (72.44%), transposable elements and repetitive sequences accounting for 33.61% and 34.99% of the genome, respectively. Among the transposable elements, long terminal repeat (LTR) sequences were predominant (30.91% of the genome), mostly composed of Copia (24.51%) and Gyspy (6.40%) LTRs.

### Gene family contraction and expansion

The comparisons between the genomes of *L. rigidum, A. thaliana*, and the four grass crop species are summarised in Figure 2. We confirmed that *L. rigidum* is very closely related to *L. perenne* (Figure 2 panel a). The four grass species, *L. rigidum, L. perenne, O. sativa* and *Z. mays* have more shared gene families than species-specific gene families, and *L. rigidum* shares more gene families with *L. perenne* than with *O. sativa* and *Z. mays* (Figure 2 panel B). There are ∼14 times more expanded than contracted gene families in *L. rigidum*. This is in striking opposition to the *L. perenne* genome where there are ∼19 times more contracted than expanded gene families (Figure 2 panel A central area). Also surprisingly, the distribution of genes with multiple orthologs, unique paralogs, and single-copy orthologs in *L. rigidum* is more similar to that of *Z. mays* than *L. perenne*. The distributions of 4DTv in Figure 2 panel C show two things: first, *L. rigidum* diverged from *L. perenne* more recently than with *Z. mays*; and second, *L. rigidum* experienced a recent WGD event while *L. perenne* experienced repeated and relatively older WGD events shown by the comparatively flatter 4DTv distribution.

**Figure 2.**
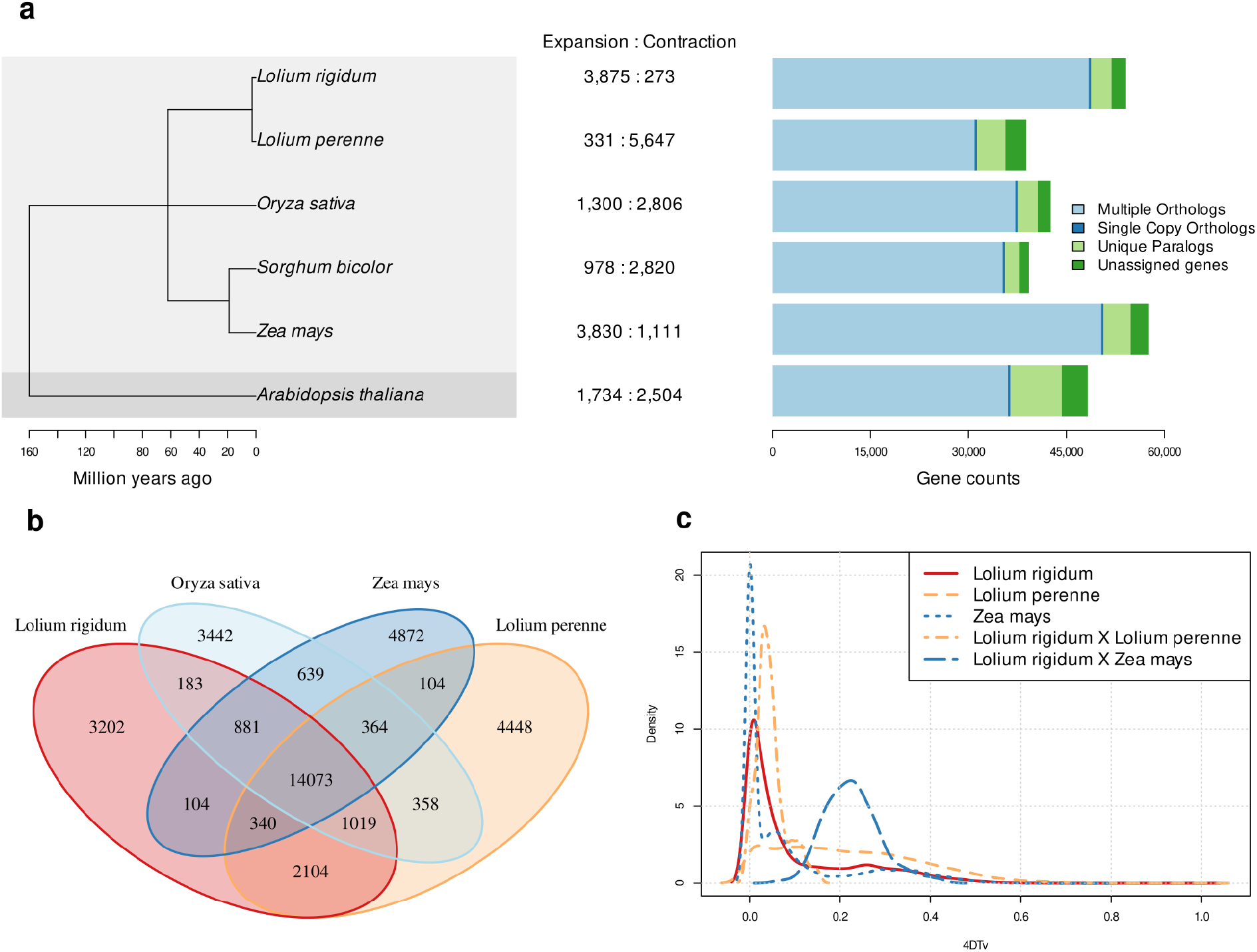
*Lolium rigidum* comparative genomics. **a (left)**: phylogeny based on single-copy gene orthologs; **a (centre)**: number of significantly expanded and contracted gene families; **a (right)**: distribution of genes with multiple orthologs, single-copy orthologs and unique orthologs, **b**: Venn diagram of shared gene families between *L. rigidum, L. perenne, O. sativa*, and *Z. mays*; **c**: distribution of the transversion rates in four-fold degenerate sites (4DTv) within orthogroups in *L. rigidum, L. perenne* and *Z. mays*.

GO term enrichment analysis of significantly expanded gene families in *L. rigidum* reveals herbicide resistance-related biological functions are significantly enriched. The top 15 significantly enriched GO terms are presented in Table 2.

**Table 2.** List of the top 15 significantly enriched gene ontology terms from the significantly expanded *Lolium rigidum* gene families.

| Rank | Biological Process | GO ID | Fold enrichment (p-value) |
| --- | --- | --- | --- |
| 1 | xylan biosynthetic process | 0045492 | 2.89 ($1.20 \times 10^{-2}$) |
| 2 | xenobiotic detoxification by transmembrane export across the plasma membrane | 1990961 | 2.81 ($2.08 \times 10^{-2}$) |
| 3 | xenobiotic export from cell | 0046618 | 2.81 ($2.08 \times 10^{-2}$) |
| 4 | lignin metabolic process | 0009808 | 2.78 ($4.28 \times 10^{-4}$) |
| 5 | xyloglucan metabolic process | 0010411 | 2.78 ($8.30 \times 10^{-4}$) |
| 6 | cell wall polysaccharide biosynthetic process | 0070592 | 2.77 ($5.36 \times 10^{-6}$) |
| 7 | hydrogen peroxide catabolic process | 0042744 | 2.75 ($2.55 \times 10^{-13}$) |
| 8 | hydrogen peroxide metabolic process | 0042743 | 2.75 ($2.55 \times 10^{-13}$) |
| 9 | cellulose metabolic process | 0030243 | 2.73 ($1.25 \times 10^{-5}$) |
| 10 | xenobiotic transport | 0042908 | 2.71 ($3.90 \times 10^{-3}$) |
| 11 | cellular component macromolecule biosynthetic process | 0070589 | 2.66 ($1.51 \times 10^{-5}$) |
| 12 | cell wall macromolecule biosynthetic process | 0044038 | 2.66 ($1.51 \times 10^{-5}$) |
| 13 | reactive oxygen species metabolic process | 0072593 | 2.65 ($1.74 \times 10^{-13}$) |
| 14 | cellulose biosynthetic process | 0030244 | 2.64 ($2.58 \times 10^{-2}$) |
| 15 | cellular response to ethylene stimulus | 0071369 | 2.62 ($4.55 \times 10^{-4}$) |

### Herbicide resistance genes

There is statistically significant evidence for the expansion of the detoxification genes tested, except for monodehydroascorbate reductase (MDAR). Interestingly, there was no evidence for significant expansion of the EPSPS gene family; however, there is some evidence for positive selection within one EPSPS gene between 556-570 bp of the consensus CDS (refer to Supplementary Figure 2). This position is conserved between *L. rigidum*, and *L. perenne*, but not with *A. thaliana* and *S. bicolor*. This putative target-site mutation in addition to the expansion of detoxification genes, hints at the possible basis of glyphosate resistance in the reference genotype.

## Discussion

### Genome assembly

A combination of long- and short-read sequencing in addition to Hi-C scaffolding proved to be sufficient to assemble a high-quality chromosome-level 2.4-Gb reference genome of *L. rigidum*. The very high proportion of duplicated genes is not uncommon in plant genomes (Panchy et al, 2016). The genome was relatively unambiguous with no evidence of polyploidisation according to the distribution of 4DTv. The simple diploid nature of annual ryegrass made it easier to resolve contig placements compared with polyploid species (Kyriakidou et al, 2018). The reference genome is mostly repetitive, consisting of long terminal repeat (LTR) families. Such expansion of LTRs in genomes has been linked to crop domestication (Qin et al, 2014; Huang et al, 2017) which annual ryegrass has been, before being predominantly classified as a noxious weed.

Despite the high-quality of this genome assembly, additional sequencing efforts can be made to improve it further. Individual chromosome sequencing can be performed, in addition to the assembly of the mitochondria and chloroplast genomes. This will consolidate the scaffolds excluded from the 7 chromosomes.

### Comparative genomics and herbicide resistance genes

The phylogeny inferred using our assembly and the reference genomes of six other plants matched the expected relationships and divergence times. *L. rigidum* expectedly diverged later from and shares more gene families with the other grass species than with *A. thaliana*. Despite being closely related and having similar genome sizes, the patterns of gene expansion and contraction in *L. rigidum* and *L. perenne* are the opposite of each other. This suggests that *L. rigidum* underwent recent single-gene duplication events which is further supported by the distribution of 4DTv. These single-gene duplication events may have been mediated by tandem duplication (gene duplication resulting in multiple paralogous genes adjacent to each other) which is supported by the proximity of the expanded detoxification genes.

The expansion of herbicide resistance-related gene families is another interesting finding, with six out of the seven detoxification genes families showing significant expansion. This, in conjunction with evidence for positive selection in one of the EPSPS genes without expansion of the whole family, suggests that the mechanism of glyphosate resistance in this specific plant is multifactorial. Glyphosate resistance here is likely achieved through a combination of intensified neutralisation of reactive oxygen species (ROS) by the increased number of detoxification enzymes, and possibly by rendering the EPSPS enzyme resistant to disruption by glyphosate molecules. Given that we have stronger evidence for the former rather than the latter, suggests that ROS scavenging by detoxification genes may be more important than preventing the disruption of EPSPS activity in aromatic amino acid synthesis in the reference genotype sequenced.

## Conclusion

We have assembled the first reference genome of the agriculturally important and noxious weed species, *Lolium rigidum*, at a high-quality and at chromosome level. This reference genome is pivotal in deciphering the genetic bases of new and emerging herbicide resistances, and the development of modern molecular tools for the management of this highly herbicide-resistant weed species. Upon analysing this reference genome representing only a single genotype, we were able to gather some evidence for the multifactorial bases of glyphosate resistance, i.e. target site resistance conferred by single point mutations within the gene, and non-target site resistance through the extensive duplication of detoxification genes. Hence, it is doubtless that this reference genome will be indispensable in future herbicide resistance mapping efforts making use of more genotypes in more sophisticated experimental designs. It will also be instrumental in the development of new and novel genomically informed weed and herbicide resistance control strategies including genomic prediction models which will improve the speed and cost-effectiveness of herbicide resistance assays.

## Data Availability

Raw Illumina, MinION, PromethION, and Hi-C reads are available under the NCBI Bioproject PRJNA799061. The reference genome assembly and the genome annotations can be found in NCBI’s database at https://www.ncbi.nlm.nih.gov/data-hub/genome/GCF_022539505.1.

The comparative genomics, plotting, and statistical analyses pipelines are described in the github repository: https://github.com/jeffersonfparil/Lolium_rigidum_genome_assembly_and_annotation.

## Author Contributions

AFL, JP, and EBB conceived the project and designed the experiments. EBB performed the nucleic acid extractions and short-read sequencing. EBB and JP performed the MinION sequencing. LC performed the PromethION and Hi-C sequencing. GP and RVR assembled the genome, submitted the assembly to NCBI, requested for the annotation, and deposited the raw data to NCBI. JP performed the comparative genomics and analysed the assembly and annotations. JP drafted the manuscript. All authors edited and contributed to the article.

## Funding

Funding for this study was partially provided by the Department of Agriculture, Water and the Environment (DAWE) and the Grains Research and Development Corporation (GRDC) (grant ID: 4-FY9JQPE), Australian Research Data Commons (grant ID: DP727) as well as by the Commonwealth Scientific and Industrial Research Organisation (CSIRO) (Applied Genomics Initiative).

## Conflict of Interest Statement

The authors declare that they have no conflicting interest.

## Acknowledgments

We thank DAWE, GRDC, ARDC and CSIRO for the funding, and the University of Melbourne for hosting this study.

## Notes

### Competing Interest Statement

The authors have declared no competing interest.

