## Supplemental Figures for "Rounding up the annual ryegrass genome: high-quality reference genome of *Lolium rigidum*"

### Supplementary Figures

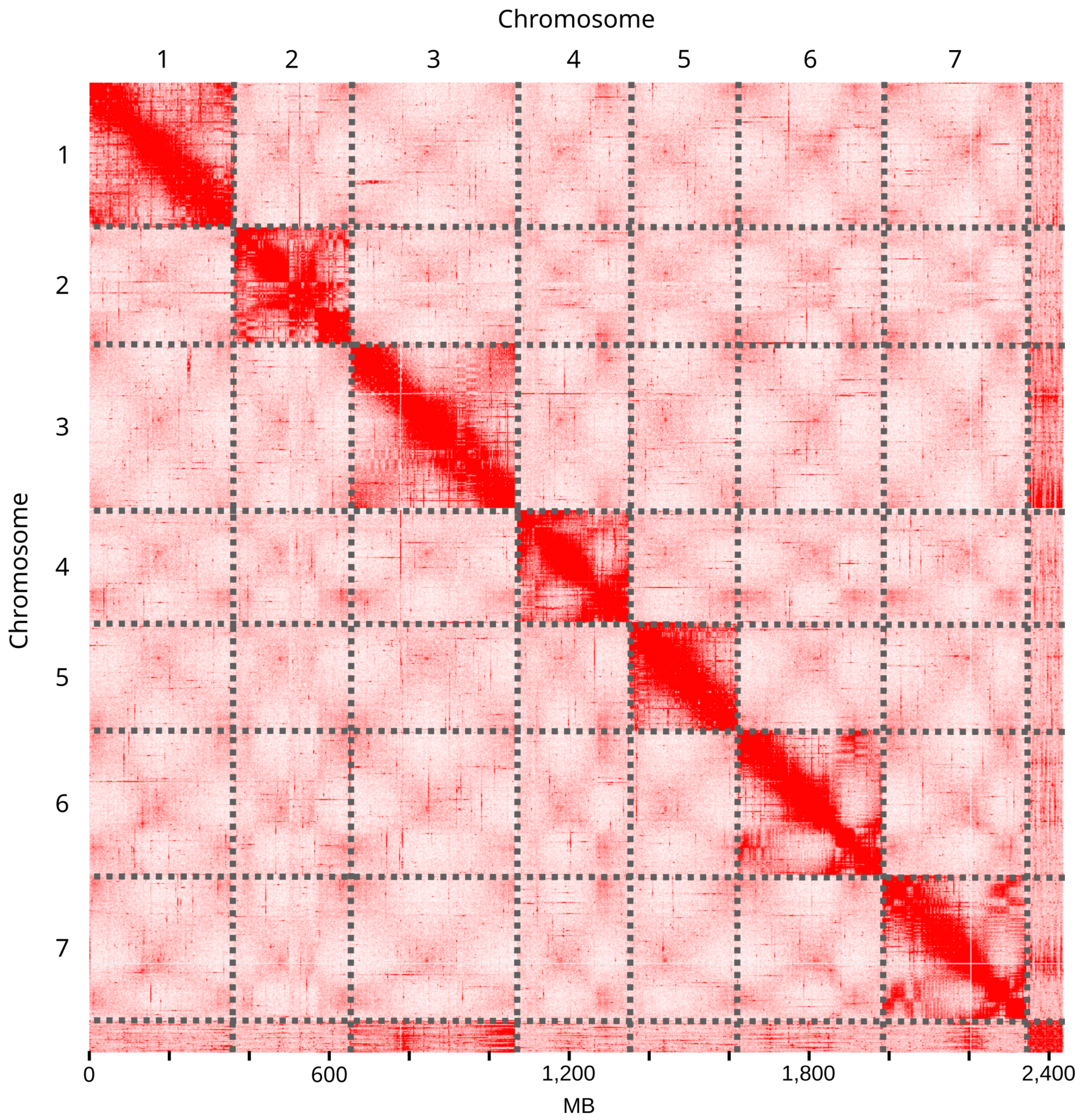


**Supplementary Figure 1**. Hi-C interaction heatmap highlighting the 7 *Lolium rigidum* chromosomes.

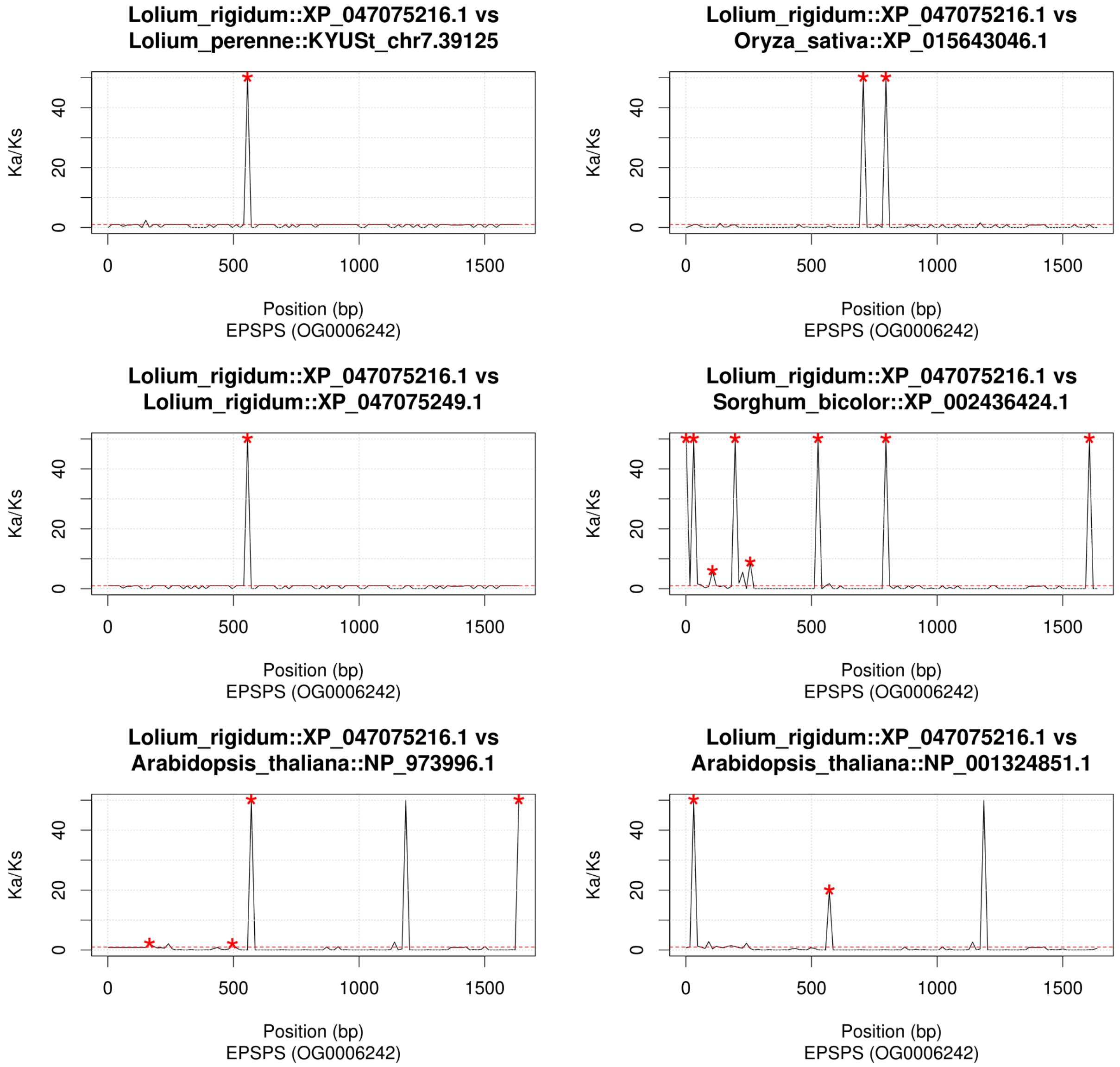


**Supplementary Figure 2**. Ka/Ks (ratio of the number of nonsynonymous substitutions per non-synonymous site to the number of synonymous substitutions per synonymous site per unit time) across non-overlapping 15-bp sliding windows comparing an enolpyruvylshikimate phosphate synthase (EPSPS) gene of *L. rigidum* (i.e. XP_047075216.1) to the EPSPS genes of *O. sativa*, *S. bicolor*, a paralog in *L. rigidum*, and two homologs in *A. thaliana*. Red asterisks show significant peaks at p≤0.001.
